# Drosophila embryos spatially sort their nutrient stores to facilitate their utilization

**DOI:** 10.1101/2022.05.20.492771

**Authors:** Marcus D. Kilwein, Matthew R. Johnson, Jonathon M. Thomalla, Anthony Mahowald, Michael A. Welte

## Abstract

Animal embryos are provisioned by their mothers with a diverse nutrient supply critical for development. In Drosophila, the three most abundant nutrients (triglycerides, proteins, and glycogen) are sequestered in distinct storage structures, lipid droplets (LDs), yolk vesicles (YVs) and glycogen granules (GGs). Using transmission electron microscopy as well as live and fixed-sample fluorescence imaging, we find that all three storage structures are dispersed throughout the egg but are then spatially allocated to distinct tissues by gastrulation: LDs largely to the peripheral epithelium, YVs and GGs to the central yolk cell. To confound the embryo’s ability to sort its nutrients, we employ mutants in Jabba and Mauve to generate LD:GG or LD:YV compound structures. In these mutants, LDs are missorted to the yolk cell and their turnover is delayed. Our observations demonstrate dramatic spatial nutrient sorting in early embryos and provide the first evidence for its functional importance.

## Introduction

After fertilization, animal embryos develop for extended periods without access to external nutrients. In oviparous species, for example, the young animal gains access to additional nutrients only after hatching when it can feed independently. Therefore, mothers provision their embryos with nutrient reserves to enable embryonic development. In Drosophila, these reserves include neutral lipids (triglycerides and sterol esters), proteins (yolk proteins) and carbohydrates (predominantly glycogen)^1-4^. These nutrients provide both energy supplies and carbon backbones for anabolic metabolism. Different nutrients are not present freely in the cytoplasm, but are segregated from each other and packaged into distinct structures^5^. Neutral lipids are present in the center of lipid droplets (LDs), ubiquitous cellular organelles in which a single phospholipid layer surrounds a central hydrophobic core. Yolk proteins reside inside membranous yolk vesicles (YVs), oocyte- and embryo-specific LROs (lysosome-related organelles) delimited by a phospholipid bilayer. Glycogen forms so-called β particles (carbohydrate chains attached to the priming protein Glycogenin) that assemble into larger glycogen granules (GGs or α particles). These nutrients are utilized at different rates and at different embryonic stages^2,3,6,7^. Most previous studies relied on biochemical analysis of whole embryos and thus could not address the spatial organization of these nutrients in the embryo. In this paper, we address this issue and find that GGs, LDs, and YVs undergo dramatic sorting in the first few hours of embryogenesis and that this sorting is a prerequisite for proper nutrient utilization.

Drosophila embryos are a syncytium for the first ∼2.5hrs. During this period, the nuclei divide 13 times near synchronously before undergoing bulk cytokinesis. The first 8 divisions (stage 1 and 2) occur deep within the embryo, with a minority of nuclei staying in the interior and most migrating to the periphery. Arrival of nuclei at the surface and formation of pole cells/germline (stage 3) mark the beginning of the syncytial blastoderm. Over the next hour (stage 4), the peripheral nuclei undergo the 9^th^-13^th^ divisions^8^. Stage 5 encompasses a 1hr long interphase where a simultaneous cytokinesis (cellularization) generates ∼6000 diploid cells organized as an epithelium at the embryo’s periphery and one central, syncytial yolk cell. The epithelium gives rise to all the tissues of the future larva and adult, while the yolk cell is a transient tissue that functions as a yolk protein depot as well as a positional cue during the maturation of ecto- and endoderm^8,9^. Gastrulation movements (stage 6) and germ-band extension (stages 7-10) then transform this 2D epithelial layer into a complex 3D body plan.

The embryo initially shows little or no spatial segregation of nutrients, with LDs and YVs homogeneously distributed^10^. In larvae and adults, in contrast, nutrients are unevenly distributed, with specialized storage tissues dedicated to receiving and disseminating specific nutrients (*e*.*g*., fat body tissue for fat storage). Nutritional specialization is already evident at gastrulation when LDs and YVs are allocated to distinct cells: most LDs to the peripheral epithelium; YVs exclusively to the yolk cell^11^. This distribution persists for the rest of embryogenesis (Fig. 1a-c). Thus, one of the first organizational events in Drosophila embryogenesis differentially sorts nutrients, foreshadowing nutrient handling by specialized tissues in later life stages.

**Figure 1.**
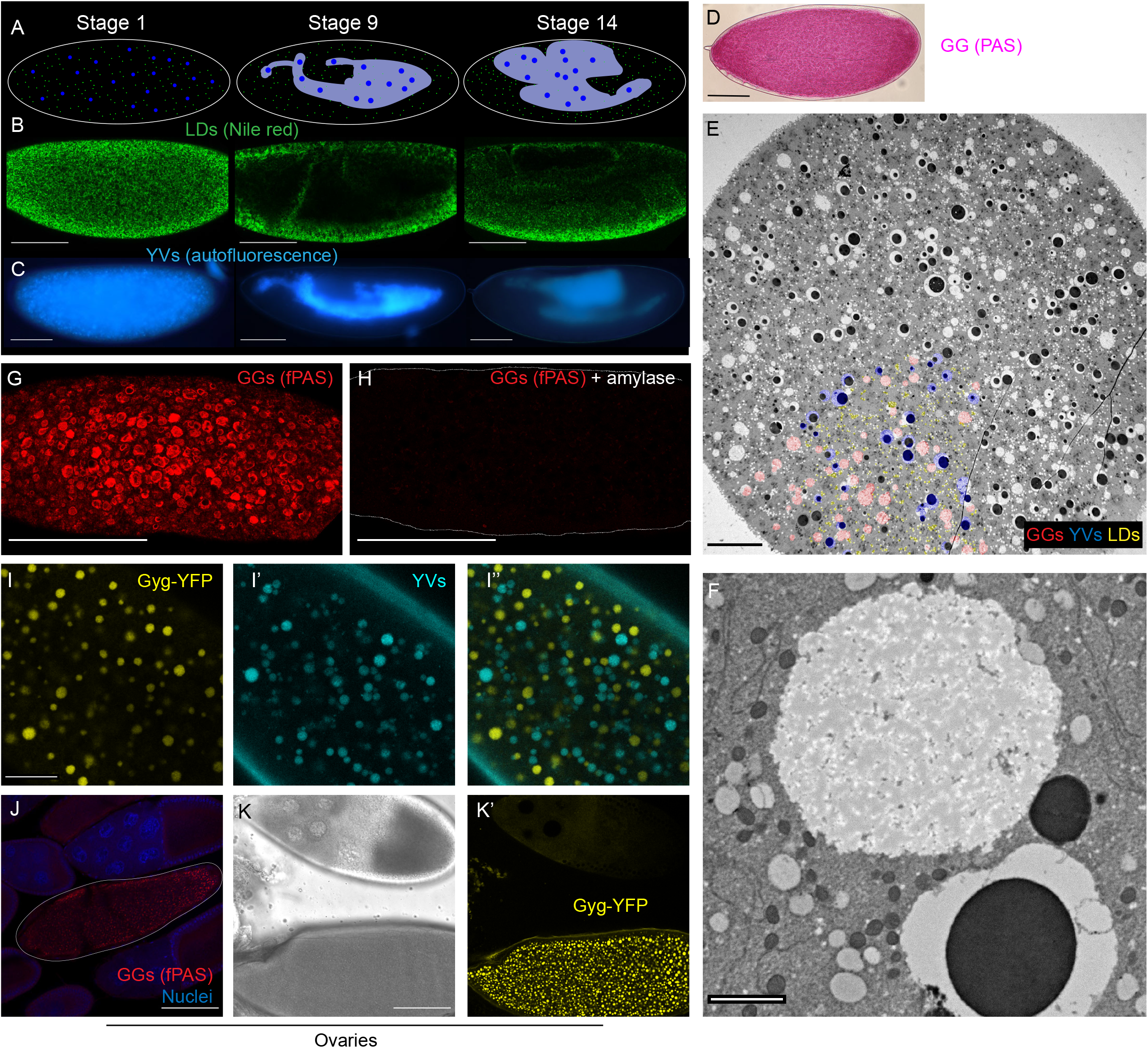
Visualizing embryonic nutrient stores. A,B,C) Distribution of LDs and YVs at three embryonic stages (stage 1 = newly laid; stage ∼ 4.5 hr old; stage ∼ 11-hr old). A) Cartoon summary, LDs are in green, YVs in blue, yolk cell in gray. B) Fixed embryos stained with Nile Red to label LDs and imaged with confocal microscopy. Scale bar = 100 µm. C) Living embryos imaged by epifluorescence microscopy to reveal the distribution of the autofluorescent YVs. Scale bar = 100 µm. D) fPAS staining of newly laid embryo visualized by brightfield. Scale bar = 100 µm. E) Cross-section of a stage 1 embryo imaged by TEM; LDs, GGs, and YVs are evenly distributed. A portion of the storage organelles are pseudo-colored, to orient the viewer, in the lower left: YVs blue, LDs yellow, GGs red. Scale bar = 20 µm. F) TEM image of a GG, the large white circle top/middle of the image. Scale bar = 2 µm. G,H) fPAS staining of newly laid embryos visualized by confocal microscopy; the embryo in H was pretreated with α-amylase for 2hrs to digest glycogen. Scale bars = 100 µm. I, I’, I’’) Confocal micrographs of a live stage 2 embryo expressing Glycogenin-YFP. Scale bar = 20 µm. I = Glycogenin-YFP, I’ = yolk autofluorescence, I’’ = merged. J,K) Glycogen accumulation in ovary follicles; scale bar = 100 µm. J) PAS and DAPI stained follicles, PAS signal is visible only in the stage 13 oocyte (outlined), with no signal in the stage 10 follicle above it). K, K’) Follicles expressing Glycogenin-YFP. K = bright field; K’ = YFP channel. GGs are not detectable at stage 10 (top) but are prominent in stage 14 (bottom).

How embryos spatially control the third major nutrient, glycogen, remains unknown. We therefore visualized GGs using Periodic Acid-Schiff (PAS) staining and fluorescence microscopy as well as a Glycogenin-YFP fusion. These approaches revealed GGs as highly dynamic, undergoing both morphological changes and redistribution before cellularization. By gastrulation, GGs are almost exclusively allocated to the yolk cell, like YVs. We also found that in embryos lacking the LD protein Jabba, LDs are tightly associated with GGs and are transported together to the yolk cell. These misallocated LDs are consumed slower than in the wild type and persist through hatching. Perturbed LD turnover is likely the result of LD misallocation, since delayed consumption is also observed in embryos in which LD misdistribution results from inappropriate interactions with YVs. We conclude that early nutrient sorting during Drosophila embryogenesis leads to an optimal nutrient allocation, ensuring that nutrients are utilized efficiently.

## Results

### Lipid droplets and yolk vesicles are sorted to different tissues by cellularization

Previous studies had suggested that LDs and YVs are likely present throughout early Drosophila embryos ^11^. However, they had relied on fluorescence microscopy of whole mount samples. To unambiguously determine the distribution of LDs and YVs, we analyzed the two organelles in cross-sections using TEM. Both LDs and YVs were homogenously distributed throughout the depth of the embryo (Fig. 1e; in the lower left, LDs are false colored in yellow, YVs in blue). Thus, early embryos start out with LDs and YVs intermixed.

In stage 2 embryos, myosin-II mediated contractions of the cortex lead to large^-^scale circular flows^12,13^. At the periphery, cytoplasm flows from the pole along the cortex to the middle of the embryo where it descends towards the interior and then flows along the long axis of the embryo to reemerge at the poles^12^ (cartooned in Supplementary Fig. 3 f). YVs are known to be carried by this flow^12^. We injected embryos with an LD-specific dye and monitored them live by confocal microscopy (Video S1). In stage 2, LDs flow in the expected pulsative manner along the periphery. In subcortical planes (Video S2), individual particles can be followed for long distances, allowing us to quantify the flow of several organelles by particle image velocimetry^14^ (see below). Because our imaging conditions do not allow us to image the middle of the embryo, we could not directly observe their flow in the embryo center. However, movies taken at 40 µm depth (Video S1) are consistent with massive rearrangement of these interior LDs and their flow towards the poles.

Previous fixed-embryo analysis had found that by Stage 3, LDs are highly enriched in the peripheral ∼40 µm of the embryo^10,15^, while YVs remain throughout. We see the same pattern in our movies, where LDs enrich at the periphery during Stage 2 and remain enriched there through Stage 5 (Video S1). During Stage 4, YVs deplete from this region^13^, and by stage 9, LDs and YVs have segregated into the epithelial cells versus the yolk cell^11^, respectively, a distribution that remains through the rest of embryogenesis (Fig. 1b,c)^10,15^.

In summary, YVs and LDs are intermixed in the early embryo and segregate from each other in two steps. During stage 2, LDs enrich in the periphery, and during stage 4, YVs deplete from the same region. As a result, the two nutrient stores are allocated to distinct cells by cellularization, creating an early nutrient differentiation between the cells of the embryo (Fig. 1a).

### Two novel, subcellular approaches to visualize glycogen granules by light microscopy

Does the third nutrient store (GGs) also undergo sorting? By electron microscopy, GGs appear as large, weakly staining, membrane-less structures (Fig. 1f). TEM cross sections of young embryos show that GGs are evenly distributed early on, like LDs and YVs (Fig. 1e, GGs in red in the lower left). Thus, all three structures are present throughout the embryo at the start of development.

To visualize GGs by light microscopy, we employed Periodic acid-Schiff (PAS) staining, a histological stain for visualizing carbohydrates. This approach works well at the organismal and tissue levels, but it does not reveal the subcellular organization into GGs. PAS-stained early embryos show signal throughout the cytoplasm, but conventional imaging approaches do not reveal fine structure (Fig. 1d). However, when adapted for fluorescence microscopy (Thomas Kornberg, personal communication), fluorescent PAS (fPAS) signal revealed discrete granular structures of <1-5µm diameter throughout the cytoplasm (Fig. 1g). When embryos were pretreated with α-amylase to degrade glycogen, fPAS signal was largely abolished (Fig. 1h), confirming that most fPAS signal in the early embryo represents glycogen. As an independent test, we analyzed fPAS signal during oogenesis, where glycogen specifically accumulates in late-stage oocytes (Stage 13 and 14)^5,16^. fPAS signal recapitulates this pattern: only the oldest oocytes were fPAS positive (Fig. 1j). We conclude that fluorescent imaging with PAS staining labels glycogen, with sufficient contrast to resolve individual granules.

Live observation of YVs and LDs has revealed detailed information about their mechanism of motion ^11^. Because β particles, the subunits of GGs, contain the priming protein Glycogenin at their center ^17^, we employed a protein trap line^18^ in which an additional coding YFP exon is inserted into the endogenous *Glycogenin* (*Gyg*) gene. By confocal microscopy, we observed structures ∼1-5 µm in diameter in stage 2 embryos of this strain (Fig. 1i). Because YFP fluorescence is destroyed by PAS staining, we could not directly compare PAS-stained GGs and Glycogenin-YFP, but several lines of evidence strongly support that Glycogenin-YFP indeed reveals GGs. First, size, distribution, and abundance of the YFP-positive structures fit the GGs visualized by PAS staining (Fig. 1g). Second, the only known abundant embryonic structures in this size range are YVs and GGs, and live imaging of embryos revealed that the autofluorescent YVs are distinct from the YFP structures (Fig. 1i,i’,i’’). Third, during oogenesis, YFP structures become distinct only in Stage 13 and Stage 14 oocytes (Fig. 1k,k’), just like GGs (Fig. 1j). Fourth, in certain mutants (see below), LDs are arranged around GGs (as observed by TEM or fPAS), and we observe the same association around Glycogenin-YFP granules (Fig. 3g’). Thus, Glycogenin-YFP reveals GGs.

As a final test, we performed *in-vivo* embryo centrifugation, a technique in which living syncytial embryos are centrifuged to separate their components by density^19,20^. GGs as revealed by PAS staining represent the densest fraction, opposite the low-density neutral lipid (Supplementary Fig. 1b); this fraction appears clear in bright light (Supplementary Fig. 1a), fitting a lipid free fraction. Similarly, in centrifuged Glycogenin-YFP embryos, YFP signal accumulates at the very bottom of the embryos, even below the autofluorescent YVs (Supplementary Fig. 1b). In centrifuged oocytes, coalesced GGs also form a cap below tightly packed YVs^5^.

### Glycogen distribution changes during development leading to yolk cell allocation

By TEM, GGs are present throughout the embryo in stages 1 and 2, just like YVs and LDs (Fig. 1e). To determine whether GGs are also spatially sorted during embryogenesis, we performed fPAS staining on embryos of different ages. In stage 1, glycogen was evenly distributed (Fig. 1G). By stage 3, imaging in a subcortical plane shows a reduction in the number of GGs at the embryo’s surface (compare Fig. 2a to Fig. 1g). In stage 5, fPAS signal was absent from the periphery; the embryo shown in Fig. 2B was imaged 40 µm below the surface (embryo is outlined in white). This developmental time course suggests that GGs deplete from the peripheral regions.

**Figure 2.**
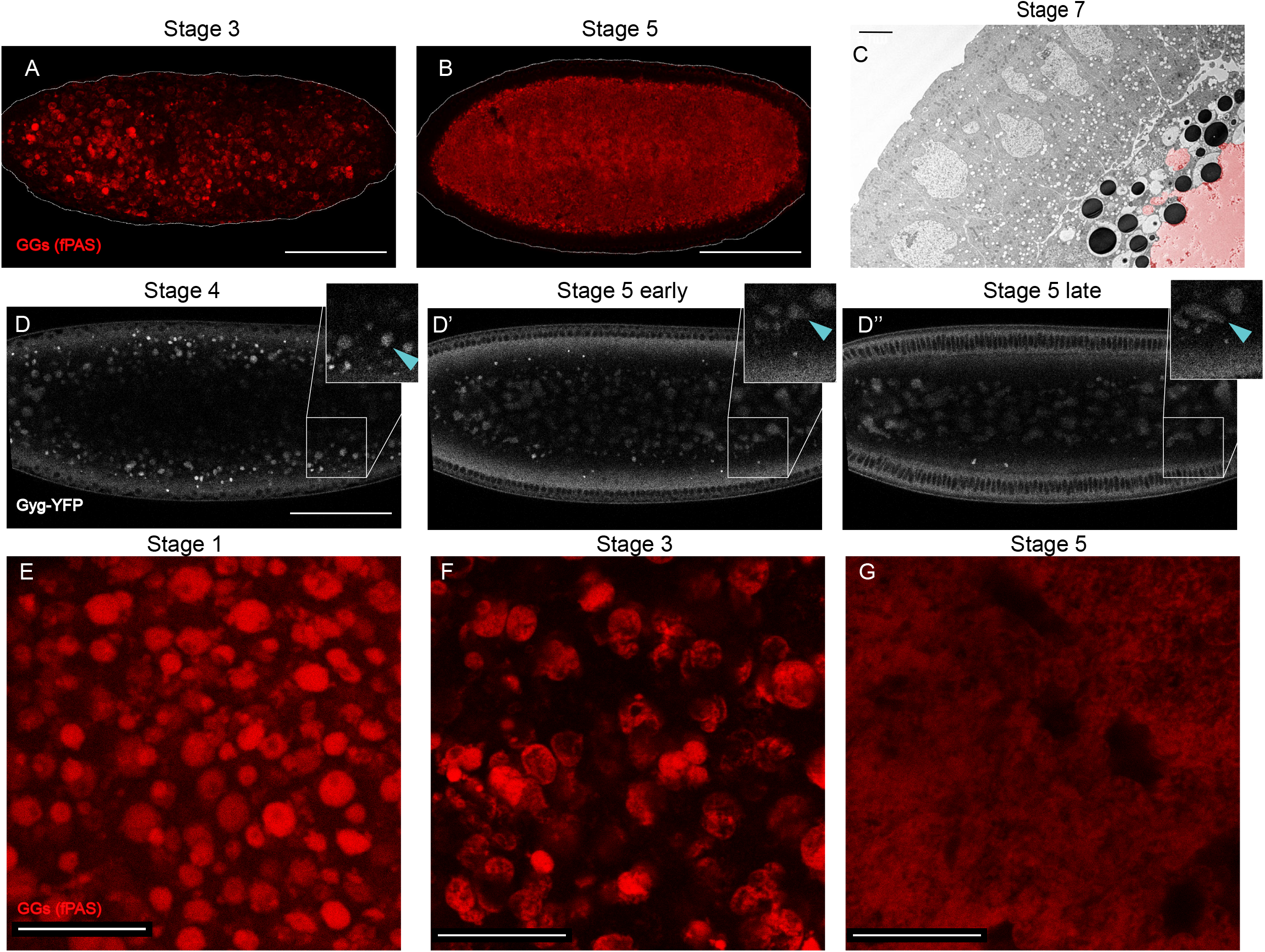
GGs change their morphology and distribution during the first 3 hours of embryogenesis. A,B) PAS staining of whole embryos, imaged by confocal microscopy. Scale bars = 100 µm. A) Stage 3 embryo with GGs beginning to coalesce. B) Stage 5 embryo with coalesced GGs localized to the embryo interior. C) TEM image of a stage 6 embryo; coalesced glycogen (false colored red) evident in the yolk cell which occupies the bottom right corner, scale bar = 5 µm. D) Frames from confocal time-lapse movie of Glycogenin-YFP expressing embryo (Video S4). Note the inward movement of the GGs and their coalescence. Scale bar = 100 µm. E, F, G) fPAS staining of embryos of different stages; scale bar = 20 µm. The reduced signal in (G) is likely a result of the deeper focal plane due to inward motion of the GGs.

Live imaging of Glycogenin-YFP expressing embryos identified discrete, mobile GGs in young embryos (Video S2). Fig.2 d-d” shows still images from a movie (Video S3) for an embryo imaged ∼40 µm below the surface through stages 4 to 5. Initially (Fig. 2d), GGs were enriched in a broad area just below the peripheral nuclei (black circles). Over time, GGs progressively moved into the interior. By the onset of stage 5 (Fig. 2d’), GGs were largely absent from this area and most were in the center. Towards the end of stage 5, the GGs were deep within the embryo’s interior (Fig. 2d’’), well below the forming cells. This inward movement was confirmed using epifluorescence microscopy (Supplementary Fig. 2a).

Thus, both fPAS and live imaging show redistribution of GGs into the embryo’s interior during stages 3-5. This pattern suggests that after cellularization the GGs are allocated to the yolk cell and are absent from the surrounding epithelium. This distribution was confirmed by TEM imaging: the stage 7 embryo in Fig. 2c has glycogen (false colored in red) entirely in the yolk cell (see also Fig. 4b). And fPAS staining of embryos in stages 9 and 10 reveals that the majority of glycogen is indeed in the yolk cell (Supplementary Fig. 2b). Overall signal is much reduced at that time, and by stage 11 we can no longer detect appreciable fPAS signal (Supplementary Fig. 2c), suggesting that glycogen is being turned over. Glycogenin-YFP is also restricted to the yolk cell in later stages (Supplementary Fig. 2c).

### GG morphology changes during development

Our fPAS and Glycogenin-YFP time courses suggest that GG morphology changes as the embryo develops. We therefore examined fPAS-stained embryos at higher magnification. Newly laid embryos were characterized by discrete GGs evenly distributed within the focal plane (Fig. 2e). In Stage 3, GGs were arranged much less homogeneously, forming clusters: frequently, multiple GGs were in close contact with each other, and there was glycogen-free space between clusters (Fig. 2f). This pattern implies that GGs not only move inwards (*i*.*e*., perpendicular to the focal plane shown), but also within the plane towards each other. Consistent with that notion, some of the GGs are no longer round, but appear as oblong or more complicated aggregates, implying that GGs are coalescing. By stage 5, fPAS signal forms a single, largely homogeneous mass (Fig. 2g) that only retains remnants of the granular structure at its outer edges (Fig. 2c). This single GG mass occupies the center of the embryo (Fig. 2b). Consistently, TEM imaging of stage 7 embryos reveals a large, fused glycogen structure (Fig. 2c, red colored area).

We observed this same morphology shift in movies of Glycogenin-YFP embryos (Fig. 2d-d”; Video S3; Supplementary Fig. 2a). In stage 4, GGs near the periphery are largely discrete structures, while GGs deeper in the embryo tend to be larger and are misshapen, consistent with fusion. Over time, as the GGs shifted towards the embryos center, they formed larger, less spherical entities (stage 5 early, Fig. 2d’; Supplementary Fig. 2a), so that by stage 5’s end, GG signal showed many amorphous structures in the embryo’s interior (Fig. 2d’’; Supplementary Fig. 2a). The insets in Fig. 2d-d’’ track a single GG indicated with a blue arrowhead as it moved inward and closer to other GGs and then fused. In summary, GGs undergo a dramatic morphological shift as they migrate into the center of the embryo.

### Lack of the LD protein Jabba results in tight association between LDs and GGs

Our analysis shows that LDs, GGs, and YVs are initially intermixed but are segregated to distinct tissues by cellularization: while most LDs end up in the peripheral epithelium, both YVs and GGs are allocated to the interior yolk cell. The enrichment of both protein and glycogen stores in the yolk cell is consistent with the idea of the yolk cell as a metabolically supportive tissues^8,9,21^. But why LDs are allocated differently and why the embryonic nutrient supply is spatially organized at all is unknown.

As a potential inroad into this problem, we re-examined embryos lacking the LD protein Jabba, one of the most abundant proteins on embryonic LDs^22^. Such embryos have normal triglyceride content but display abnormal LD distribution in stage 4^22^. When we imaged fixed stage 1-2 embryos stained for LDs, wild-type embryos displayed the pattern expected from our TEM analysis (Fig 1e); LDs were absent from many circular areas (presumably GGs and YVs) but distributed evenly throughout the remaining spaces (Fig. 3a). The pattern in *Jabba* mutants (two different alleles, *Jabba*^*DL*^ and *Jabba*^*zl01*^) was dramatically different. Here most LDs were accumulated in rings, with few LDs occupying the space between the rings (Fig. 3b, and data not shown). This aberrant distribution was not an artefact of fixation, as we saw the same pattern in embryos injected with LD dyes and imaged live (Supplementary Fig. 3d, 0s panel).

**Figure 3.**
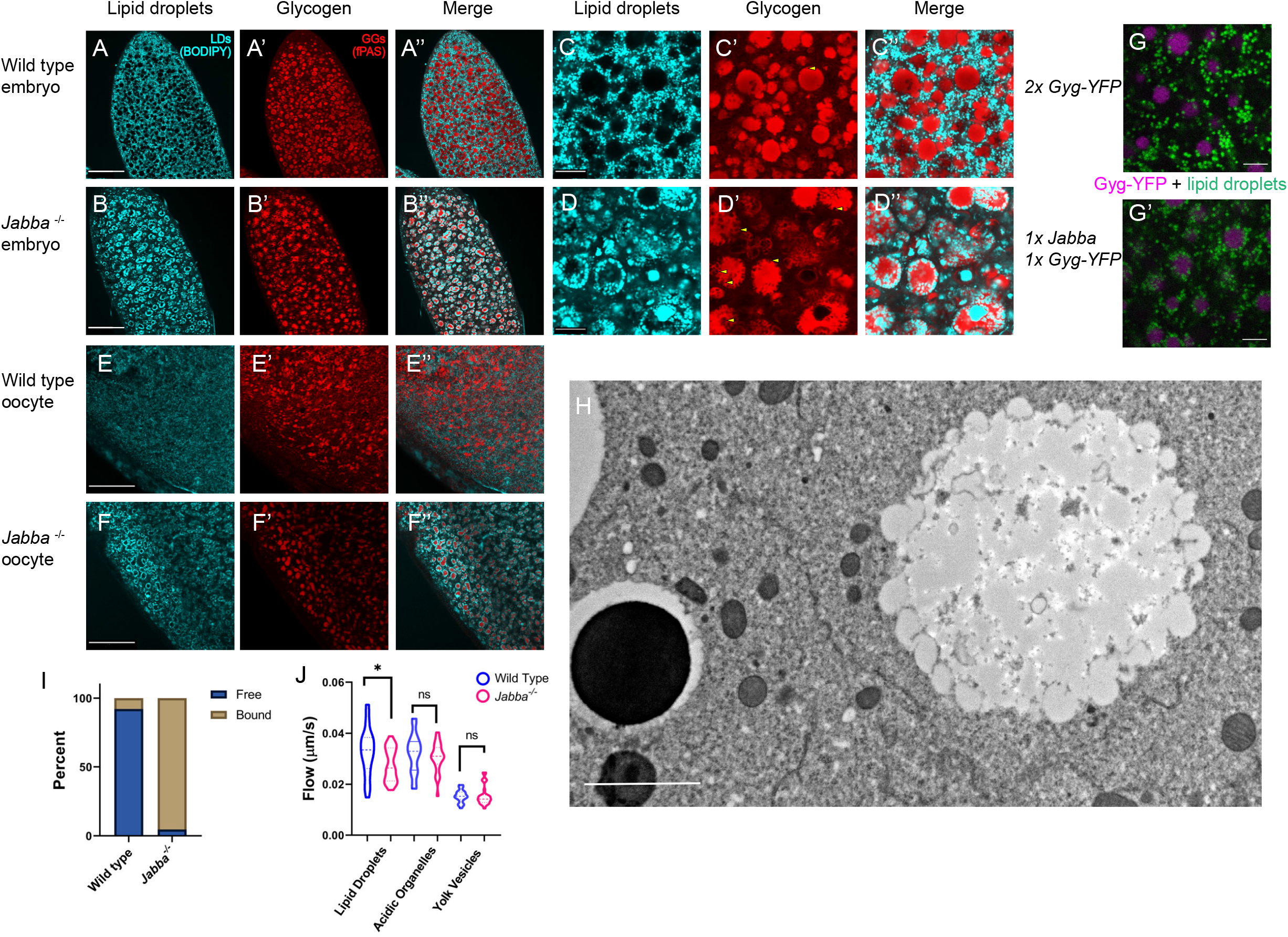
LDs and GGs aberrantly interact in *Jabba* mutant embryos. A-F’’) LDs (cyan, BODIPY) and GGs (fPAS, red). A-B’’) Stage 2 embryos; scale bars = 60 µm. A-A’’) wild type, B-B’’) *Jabba*. C-D’’) the same embryos at a higher magnification. Scale bars = 10 µm. Arrowheads (C’&D’) indicate indentations in GGs occupied by LDs. E-F’’) Late-stage oocytes stained for LDs (cyan, BODIPY) and GGs (fPAS, red). Scale bars = 40 µm. GG-LD association is already visible in F’’. G,G’) LDs (green, LipidSpot610) and GGs (magenta, Glycogenin-YFP) in Glycogenin-YFP (G) and 1 x Glycogenin-YFP,1 x *Jabba* (G’) embryos. 1x *Jabba* embryos display LD-GG association not found in the wild type. Scale bars = 5 µm. H) TEM image of *Jabba* embryo; GG with its surface decorated by bound LDs. LDs are largely absent from the cytoplasm. Scale bar = 2 µm. I) quantification of the number of LDs bound to GGs vs free in wild-type and *Jabba* embryos. Quantitation of 3 TEM images per genotype (>400 LDs) revealed that the majority of LDs are not associated with GGs in the wild type, while ∼95% of LDs are bound to GGs in *Jabba* embryos. J) PIV analysis of motion of LDs, lysosomes (acidic organelles), and YVs in wild-type and *Jabba* embryos. The variation in speeds for a given organelle results from the pulses at the embryo’s cortex. Mean LD velocity in wild type versus *Jabba*: 0.034 vs. 0.028 µm/s; p = 0.0144. Mean velocity of acidic organelles: 0.032 vs 0.031 µm/s, p = 0.406. Mean velocity of YVs: 0.015 vs 0.016 µm/s, p = 0.815. Analysis based on at least 3 embryos per genotype, with 9 measurements per embryo, i.e., n values of at least 27. p-values generated by unpaired, two tailed t-tests. * p<0.05.

**Figure 4.**
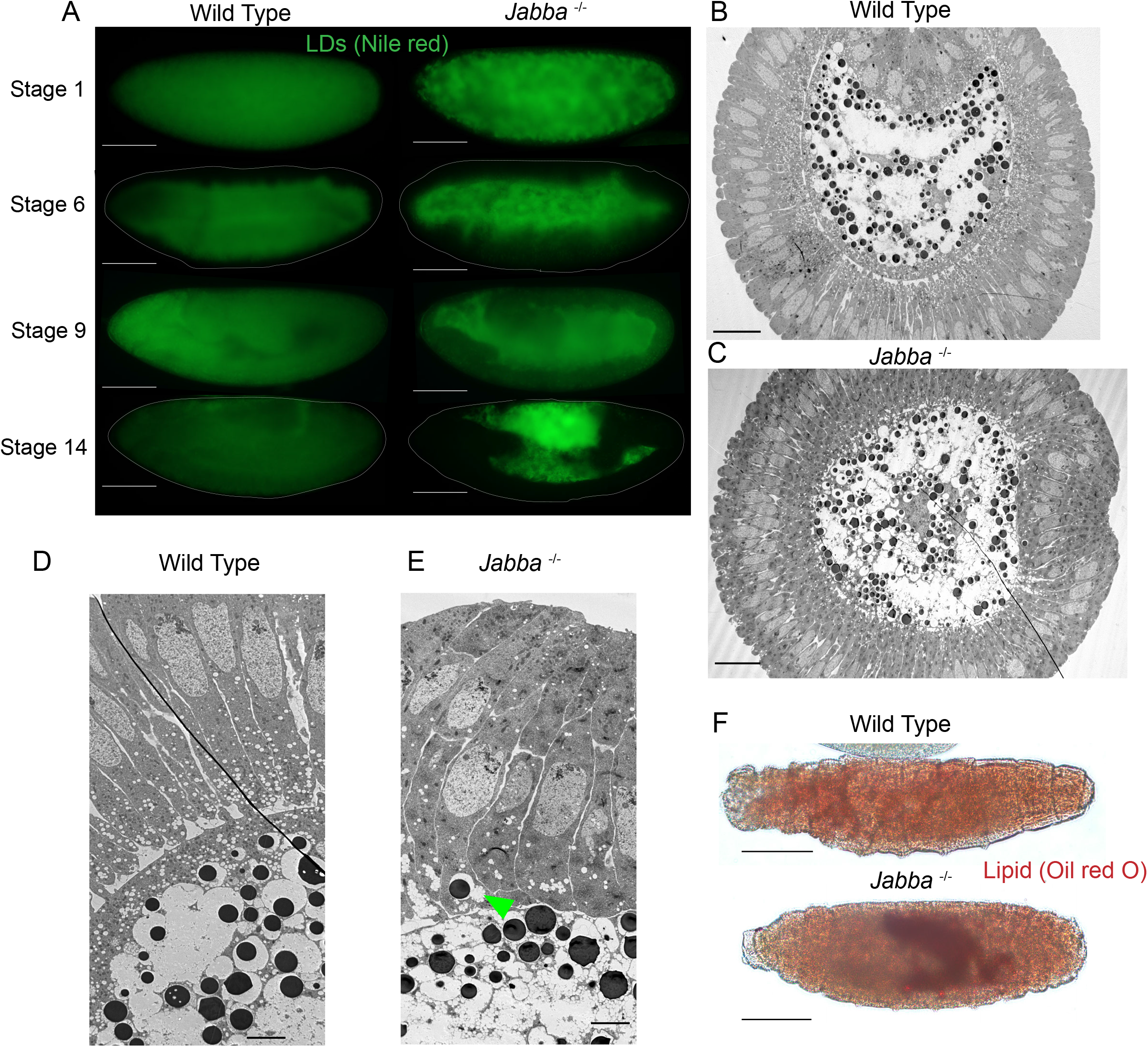
In *Jabba* mutant embryos, lipid droplets are mislocalized and their consumption is disrupted. A) Wild-type and *Jabba* embryos of various stages stained with Nile red to detect LDs and imaged with epifluorescence microscopy. Scale bars = 100 µm. B,C) TEM cross sections of stage 6 embryos. B) ventral furrow is at the top of the image. C) the ventral furrow is at the right of the image. Scale bars = 100 µm. D,E) TEMs of stage 6 embryos at higher magnification showing the border between the cellularized epithelium (right) and yolk cell (left). Scale bars = 5 µm. The green arrowhead (E) shows a YV attached to LD:GG complex which has inappropriately localized to the epithelium. F,G) L1 larva stained with Oil Red O to detect LDs. Scale bar = 80 µm. The *Jabba* larva’s residual LDs are visible in its gut.

The pattern in *Jabba* mutants suggests that LDs accumulate around YVs or GGs. Co-detection of LDs and YVs revealed no association between them (Supplementary Fig. 3e). In contrast, co-labeling of GGs and LDs showed that LDs were surrounding GGs (Fig. 3A’-B”). This finding was further confirmed in TEM cross sections (Supplementary Fig 4A). A similar, but less pronounced association was already observed when *Jabba* dosage was reduced (Supplementary Fig. 3c *1x Jabba*). We employed this observation to confirm LD-GG association in living embryos. We injected LipidSpot610 into embryos from mothers expressing Glycogenin-YFP and either one or two copies of the wild-type *Jabba* gene. LDs and GGs displayed minimal association in the otherwise wild-type background (Fig. 3g), while many LDs were present at or near the surface of GGs when *Jabba* dosage was reduced (Fig. 3g’). We conclude that when Jabba levels are reduced, LDs associate with GGs. This association can already be observed in oocytes, as soon as GGs are detected (Fig. 3e-f’). Finally, we detected no association between LDs and GGs when other LD proteins are missing, namely PLIN-2/LSD-2, Sturkopf/CG9186, or Klar (Supplementary Fig. 3c). We conclude that Jabba uniquely suppresses inappropriate interactions between these storage structures.

Although we occasionally observe LDs deep within GGs of *Jabba* mutants, for the most part the LDs are arranged in a ring pattern with glycogen in the center (Fig. 3d”). In fact, LDs appear embedded within the outer regions of the GGs; see, for example, Fig. 3d’,d”, where GGs display indentations/areas of exclusion in the PAS signal (Fig. 3d’, arrowheads). These regions are filled with LDs (Fig. 3d”). Rarely were such associations or indentations observed in wild type (Fig. 3c’, arrowheads). TEM confirmed a tight LD-GG association: in *Jabba* embryos, LDs appear to directly contact GGs, with glycogen bulging out between LDs (Fig. 3h; Supplementary Fig. 3b). In the wild type, such association is observed rarely (Fig. 1e; Supplementary Fig. 3a; Fig. 3i). This association is strong enough that when *Jabba* embryos are centrifuged, the dramatic separation of LDs and GGs into distinct layers at opposite ends of the embryo observed in the wild type (Supplementary Fig. 1b) is disrupted. In the *Jabba* mutant, glycogen signal is found intermixed with the LD layer in the region of lowest density, and pockets of LD signal are present within the glycogen layer in the highest-density region (Supplementary Fig. 1c).

### Consequences of LD:GG interactions on LD motility

To determine if the LD/GG association in *Jabba* embryos affects LD motility, we labeled LDs by injecting dyes into embryos, monitored their behavior live, and quantified their flow speeds by particle image velocimetry^14^. In the wild type, LDs and acidic organelles flow faster than YVs (Fig. 3j), presumably due to their smaller size. *Jabba* mutant embryos displayed the expected ring-arrangement of LDs (Supplementary Fig. 3d); these rings moved as a unit (Video S5), reinforcing that LD:GG complexes are stable structures. Compared to wild type, they displayed stuttered motion, minimal displacement (Supplementary Fig. 3d), and lower average velocity (Fig. 3j). In contrast, YVs and acidic organelles showed similar mobility between the two genotypes (Fig. 3j). Thus, altered LD flow in *Jabba* embryos does not represent a general defect in cytoplasmic streaming, but rather a specific disruption of LD motility, likely due to the much larger size of LD:GG complexes relative to individual LDs.

### Consequences of LD-GG interactions on LD allocation

During wild-type development, GGs and LDs are initially intermixed and homogeneously distributed throughout the embryo (Fig. 1e, Supplementary Fig. 4a). LDs enrich at the periphery by stage 3 (Video S1), and by the end of stage 5, GGs and LDs are segregated from each other and allocated to different tissues. In the absence of Jabba, GGs and LDs form large composite structures that are also distributed throughout the early embryo (Supplementary Fig. 4B). But because these composite structures travel together, segregation to distinct locations seems no longer possible. Indeed, in our movies with labelled LDs (Video S5, S6), the LDs trapped in GG:LDs complexes were far less mobile, engaging in delayed, stuttered motion and did not enrich in the periphery as in the wild type (Video S1). For post-cellularization embryos, TEM analysis revealed a massive redistribution of LDs in *Jabba* embryos relative to wild type (Fig. 4b-e), with fewer LDs in the peripheral epithelium and more in the yolk cell. In the wild type, 72% of 510 LDs scored were present in the epithelial cells, while in *Jabba* embryos it was only 23% of the 696 LDs scored.

In contrast, we only found minor differences in glycogen distribution between the two genotypes. By fPAS staining, wild type and *Jabba* were very similar (Supplementary Fig. 2E). By TEM, the bulk of GGs in *Jabba* embryos were appropriately localized to the yolk cell, just like in the wild type (Fig. 4B-E). There were a few instances of small LD:GG complexes segregated into blastoderm cells (Fig. 4E green arrowhead). We conclude that the composite LD-GG structures in *Jabba* embryos are allocated like GGs, leading to LD mislocalization.

As an independent approach, we detected LDs by Nile red staining and fluorescence microscopy in whole-mount embryos (Fig. 4a). Already in early stages, LD distribution looks different: signal is diffuse throughout the embryo for wild type and granular in *Jabba* mutants, presumably reflecting LD enrichment around GGs. At the beginning of gastrulation, signal in the mutant is more prominent in the yolk cell, a pattern that becomes even more pronounced in later stages. By stage 14, LD signal is absent from the yolk cell and present everywhere else, while in *Jabba* embryos this pattern is reversed (Fig. 4A). We conclude that in *Jabba* embryos the bulk of LDs are indeed mislocalized to the yolk cell.

In late-stage embryos, LD signal in *Jabba* embryos was not only restricted to the yolk cell but also displayed increased intensity. This difference even persisted post-hatching. LD staining using Oil Red O revealed strong signal in the gut lumen (the location of the yolk cell remnant) of newly hatched *Jabba* larvae (Fig. 4f), unlike wild type (Fig. 4f). Thus, *Jabba* mutants not only mislocalize their LDs, but fail to consume them properly.

### LD interactions with YVs also lead to their mislocalization and persistence

LD consumption in the mutant might be impaired because Jabba is important for LD breakdown or because the yolk cell is not equipped to metabolize this high concentration of LDs. To distinguish between these possibilities, we took advantage of a recent report that embryos lacking the Mauve protein display an interaction between LDs and YVs^23^. Mauve is a resident protein on lysosome related organelles (LROs) important for their maturation. YVs are a type of LRO and in the absence of Mauve display several phenotypes, including an association with LDs ^23^. Might these YV-LD interactions affect nutrient sorting?

As strong disruption of Mauve severely impairs early embryonic development^23^, we generated mothers transheterozygous for two weak alleles, *mauve*^*Rosario*^ and *mauve*^*3*^. These embryos displayed LDs association with YVs (Fig. 5a arrow), as well as the YV size heterogeneity and autofluorescence variability reported for stronger allele combinations (Fig. 5a)^23^, but no obvious association between GGs and LDs (Fig. 5b)^23^.

**Figure 5.**
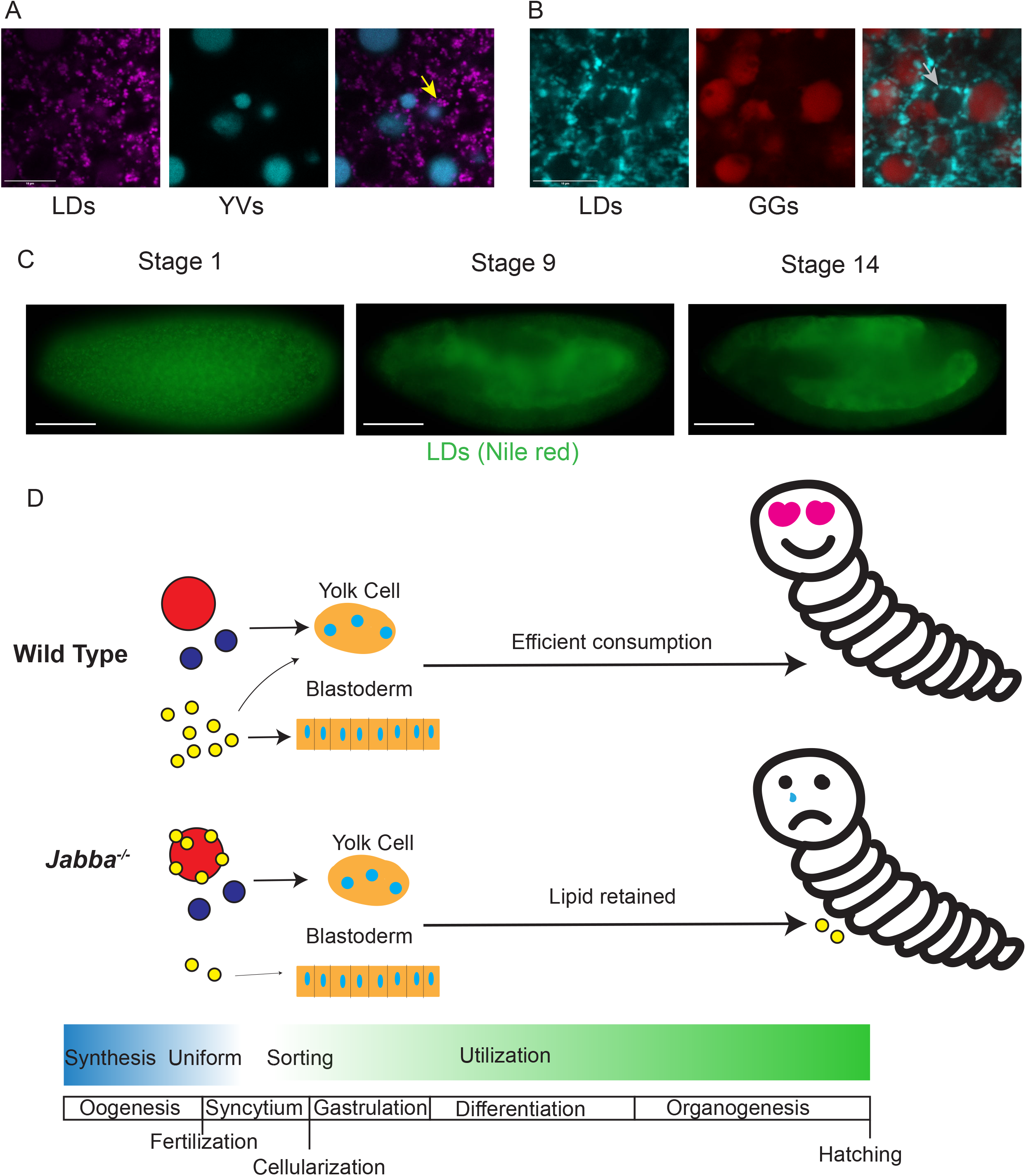
Disrupting LD localization disrupts LD consumption. A) Live imaging of LDs (magenta, Lipid Spot 610) and YVs (blue, autofluorescence) in newly laid *mauve* mutant embryos. LDs are associating adjacent to YVs. Note that the autofluorescence signal may not extend to the furthest edges of the YV due to the double membrane structure of YVs. B) Fixed imaging of LDs (blue, BODIPY) and GGs (red, fPAS) in *mauve* embryos. Arrow indicates rare LD ring not associated with fPAS signal. We believe that this likely represents LDs around YVs with undetectable autofluorescence. A,B) Confocal microscopy; scale bar = 10 µm. C) LD (green, Nile red) distribution in fixed *mauve* embryos. Stage 1: LD clustering, presumably around YVs, reminiscent of GG:LD cluster seen in *Jabba* mutants. Stage 9 and 14: mislocalization of LDs to the yolk cell. Images by epifluorescence microscopy; scale bars = 100 µm. D) Model: nutrient sorting from oogenesis through embryogenesis, comparing wild type with the disruptions seen in *Jabba* mutants. Note that the timeline is not scaled to absolute developmental time but is meant to show the brief life of nutrients through synthesis, distribution and consumption when compared to other developmental events. Mislocalized LDs in *Jabba* mutants are speculated to negatively affect completion of embryogenesis or the larvae itself.

The *mauve*^*Rosario/3*^ YV-LD association was an exciting opportunity to test our model that LDs are missorted during cellularization if they are associated with a structure destined for yolk cell deposition, allowing us to utilize YVs instead of GGs. When the mutant embryos were stained for LDs (Fig. 5c), signal in stage 1 was clustered instead of diffuse like in wild type (Fig. 4a). At stage 9, LDs signal was predominately in the yolk cell, but also present in the other tissues (Fig. 5c). At stage 14, LD staining was clearly enriched in the yolk cell (Fig. 5a) relative to wild type (Fig. 4a). Thus, mutations in two unrelated genes, *Jabba* and *mauve*, show that when LDs inappropriately interact with other nutrient structures, they are mislocalized to the yolk cell and are turned over more slowly.

## Discussion

Glycogen is a major energy store in animals, uniquely capable of providing energy rapidly, both aerobically and anaerobically. While glycogen’s functions are well investigated in adult tissues, its role during embryogenesis is less understood. In this study, we developed new tools to determine the spatial distribution of glycogen in Drosophila oocytes and embryos. Consistent with previous biochemical and electron microscopic analysis, we find that glycogen stores accumulate late in oogenesis and are organized into large, membrane-less structures. These GGs are evenly distributed throughout the early embryo and undergo two types of transitions during syncytial stages. First, they are displaced from the subcortical region during stage 4, leading to their accumulation in the center and allocation to the yolk cell. Second, simultaneously, GGs fuse into larger and larger structures, so that by the end of stage 5, most glycogen is present in a large superstructure in the yolk cell. LDs and YVs also start out with an even distribution and are then specifically allocated. After stage 5, YVs are restricted to the yolk cell, like GGs, while LDs are predominately sorted to the peripheral epithelial cells. In two different mutant conditions, we find physical interactions between LDs and either GGs or YVs. In both cases, LDs are misallocated to the yolk cell. In *Jabba* mutants, a portion of these LDs fail to be consumed during embryogenesis and persist into larval stages; in *mauve* mutants persistence through embryogenesis is milder, consistent with less severe mislocalization of LDs. We conclude that mislocalizing LDs early in embryogenesis affects subsequent LD consumption.

GGs in Drosophila oocytes and embryos are akin to the α particles that mediate long-term glycogen storage in many tissues, including muscle, liver, and fat body^17,24^. Their large size (which must correspond to thousands of β particles^5^ as opposed to ∼30 in liver^24^) likely protects against glycogen breakdown and facilitates their physical movement for differential allocation. How α particles assemble and disassemble remains unclear. Proposed binding agents holding neighboring β particles together include covalent links between glycogen chains^25,26^ or Glycogenin molecules at the surface of the β particles^27^, in addition to the Glycogenin dimer in their cores. We speculate that the amorphous mass of glycogen in stage 5 embryos represents partial dissolution of GGs into β particles, as a prelude to enzymatic breakdown of glycogen.

Consistent with this notion, fPAS staining in stages 9-10 becomes very weak (Supplementary Fig. 2b,c) and biochemical measures of glycogen levels show the same drop^2^. This timing for glycogen depletion overlaps with a proposed switch in embryonic metabolism from carbohydrate-based to triglyceride-based energy production^2^. Disassembly into β particles might provide enhanced access for the cytosolic glycogen phosphorylase, responsible for most glycogen turnover in embryos^28^.

Going forward, Drosophila oocytes and embryos should be a powerful model for unraveling the mechanism of the conversion between α and β particles. GGs are large enough to be followed by light microscopy, their assembly and disassembly occurs quickly (within ∼2hrs or less), and Glycogenin-YFP allows live imaging of these processes. In fact, to our knowledge this is the first example of live imaging of glycogen in any system.

Although during oogenesis the three major nutrient stores are made at different at times and through different mechanisms^5,11^, they all start out intermixed and homogenously distributed in the early embryo. Nutrient sorting starts in stage 2 when myosin-II driven cortex contractions establish large-scale cytoplasmic flows throughout the embryo. Flow speeds, as estimated from the behavior of YVs, are sufficient to spread out the interior nuclei along the entire anterior-posterior axis^12^. As LDs flow faster than YVs (Fig. 3j), these flows should be able to transport most LDs from the center to the poles. If these LDs are somehow captured at the periphery, reducing their return to the embryo center, it explains their enrichment in the periphery by stage 3. Consistent with an important contribution from cytoplasmic flow, the LD-GG aggregates in *Jabba* mutants flow with reduced speeds and LDs fail to enrich at the embryo surface (Video S4). Thus, we propose that this cytoplasmic flow promotes the first step of nutrient sorting, when LDs accumulate peripherally.

By stage 4 and 5, the nuclei at the embryo surface set up an array of radially oriented microtubules that traverse a ∼40 µm peripheral zone^10^. These microtubules are proposed to push YVs into the interior ^13^, and GGs may be displaced by the same mechanism, as they deplete from this zone at the same time. LDs, in contrast, move bidirectionally along these microtubules, employing cytoplasmic dynein and kinesin-1^29,30^; this motion confines them to the microtubule zone^15^. We propose that MTs keep LDs in the peripheral zone, while they push GGs and YVs inward, resulting in the second sorting step. Our analysis of *Jabba* and *mauve* mutants reveals that successful sorting also requires nutrient stores stay separated. When LDs are tightly associated with either GGs or YVs, sorting fails, and LDs are misallocated to the yolk cell.

Presumably the nutrients stored in GGs, YVs, and LDs all support the metabolic needs of the developing embryo. Why then are they allocated differently? One reason might be the different properties of their breakdown products. YVs’ amino acids and GGs’ glucose are water soluble, making diffusion through gap junctions or membrane transporters viable options for dissemination. Thus, the yolk cell can serve as a hub for glucose and amino acid distribution, as it remains connected via cytoplasmic bridges to the blastoderm through stage 9^21,31^ and expresses numerous nutrient transporters. In contrast, free fatty acids (FAs) generated from LD breakdown are poorly water-soluble and potentially toxic^32,33^; they are typically immediately channeled into specific intracellular pathways^34^. For example, efficient FA transfer from LDs to mitochondria requires proximity and direct contact^35-37^. Thus, allocation of LDs predominately to the periphery would allow efficient local energy production. Intriguingly, during zebrafish embryogenesis, LDs are initially highly enriched in the future yolk sac but are imported into the embryo proper via cytoplasmic bridges and actin-myosin based motility^38,39^. Thus, the zebrafish embryo may also depend on a local LD supply to support its dividing cells.

What are the consequences of mislocalizing LDs to the yolk cell? Our data indicate that a fraction of these LDs persist through the end of embryogenesis. Although *Jabba* mutant embryos are viable^22^, their progression through embryogenesis has not yet been analyzed in detail. Recent work on embryonic glycogen metabolism suggests that even minor disruption of LD metabolism has the potential for widespread effects on embryogenesis. Embryos that either lack glycogen reserves or are unable to access them display widespread changes in their metabolome as well as hatching delays^28^. Since fat contributes roughly 10x the energy of glucose (derived from glycogen) during Drosophila embryogenesis^3^, even the modest retention of LDs in *Jabba* mutants might have prominent effects on development.

LDs and GGs co-exist not only in embryos, but also in mature tissues (*e*.*g*., muscle, intestinal epithelia, liver, fat body)^17,40,41^. It is conceivable that in embryos inappropriate interactions between LDs and GGs are particularly harmful because of the large size of GGs and the extensive cytoplasmic streaming which presumably leads to many encounters between these organelles. It will therefore be interesting if mechanisms to keep LDs and GGs apart are specific to embryos or also important in other cells.

By devising novel imaging methods for glycogen storage structures, we have shown that Drosophila embryos dramatically reorganize their nutrients by cellularization, with distinct nutrients sorted into separate nascent tissues. The embryo employs multiple mechanisms to get its nutrient stores to the correct location, including cytoplasmic streaming, preventing inappropriate interactions, and microtubule-dependent transport. We also provide the first evidence that correct spatial allocation of LDs is necessary for their efficient consumption.

Together, these observations suggest that embryos need to achieve an optimal nutrient allocation to support subsequent steps in development (cartooned in Fig 5d) and that the spatial allocation of nutrients is essential to fully understand embryonic metabolism. The importance of this allocation is particularly remarkable as these nutrient stores only exist transiently and are consumed by the end of embryogenesis.

## Methods

### Origin of fly strains

Oregon R was used as the wild-type strain. *Jabba*^*DL*^ and *Jabba*^*zl01*^ were generated previously in the lab and are strong loss-of-function alleles with no Jabba protein detected in early embryos^22^. *Df(2R)Exel7158*/*CyO* carries a large deletion that encompasses *Jabba* and is used to reduce *Jabba* dosage; for simplicity, embryos from mothers carrying this deletion are referred to as *1x Jabba*. This stock was obtained from the Bloomington *Drosophila* Stock Center (BDSC: 7895; FLYB: FBab0038053)]. The hypomorphic alleles *mauve*^*3*^ and *mauve*^*Rosario*^ (described in^23^) were a gift from Ramona Lattao. The YFP insertion in the Glycogenin locus was generated in a large genetic screen^18^ and was obtained from the Kyoto stock center (DGRC # 115562).

### Microscopy

Laser scanning confocal microscopy was performed on a Leica SP5 equipped with HyD detectors, using either a 40x objective to show most of the embryo, or a 63x objective for subregions. Epifluorescence imaging was performed on a Nikon Eclipse E600 using a 20x objective. All images were assembled in Adobe Illustrator.

Videos, except for Supplementary Video 4, were captured at 1 frame per 30 second and are displayed at 20 frames per second. Supplementary Video 4 was captured at 1 frame per 15 minutes and displayed at 1 frame per 0.8 seconds. The orientation of the embryo was chosen to maximize the amount of the embryo captured in frame. Video processing was performed in FIJI (NIH).

Sample sizes were determined as follows. For live imaging, at least three embryos were imaged per genotype per experiment. For fixed samples, the stainings were performed at least twice. For TEM analysis, the core facility was given ten appropriately staged embryos per genotype per experiment, and then chose which were imaged based on staining success.

Exclusion criteria for imaging embryos were predetermined. Embryos not of the stage of interest, determined to have expired during preparation or image acquisition, or which were imaged in the incorrect orientation/focal depth were excluded.

### Periodic acid Schiff (PAS) and LD staining

Embryos were collected on apple juice plates for the desired time range and dechorionated with 50% bleach and fixed for 20 min using a 1:1 mixture of heptane and 4% formaldehyde in phosphate-buffered saline (PBS). To detect GGs, embryos were devitellinized using heptane/methanol and subsequently washed three times in 1xPBS/0.1% Triton X-100. Embryos were incubated first in 0.1M phosphatidic acid (pH 6) for 1hr and then in 0.15% periodic acid in dH^2^O for 15min. After one wash with dH^2^O, embryos were incubated in Schiff’s reagent (Sigma-Aldrich) until the embryos went from uncolored, to pink, to red (∼2 minutes). To stop the reaction, it was quenched with 5.6% sodium borate/0.25 normal HCl stop solution for at least 2 minutes with agitation. After replacing half the volume of the stop solution with an equal volume of 1×PBS/0.1% Triton X-100 to reintroduce detergent, the sample was shaken vigorously to free embryos stuck to the container or each other. For subsequent imaging, embryos were mounted in either Aqua-Poly/Mount, Polysciences, or glycerol (90% glycerol, 10% PBS). Mounts with ‘antifade’ or O^2^ scavenging additives should be avoided. To determine the specificity of the PAS signal, fixed and devitellinized embryos were first incubated with α-amylase (Porcine, Sigma Aldrich, 0.2mg/mL in PBS, incubated for 2hrs) to specifically digest the α-(1,4) glycosidic linkages in glycogen.

To detect LDs by staining, the methanol step needs to be omitted as it extracts neutral lipids. Instead, embryos were washed extensively with 1×PBS/0.1% Triton X-100 to remove residual heptane in a wire mesh basket then transferred to a 1.7mL microcentrifuge tube. They were then washed 2x with 1×PBS/0.1% Triton X-100. If costaining with PAS, proceed to the phosphatidic acid incubation step, adding 1µL of 1mg/mL BODIPY 493/503, Invitrogen, then proceeding with subsequent steps as normal. Red lipid dyes overlap Schiff’s reagent’s spectra and should be avoided for costaining.

For staining of lipid droplets without PAS costaining, remove the 1×PBS/0.1% Triton X-100 wash, and replace with 1×PBS/0.5% Triton X-100/10% BSA/0.02% sodium azide (toxic) and incubate for 1 hr. Replace the solution with fresh 1×PBS/0.5% Triton X-100/10% BSA/0.02% sodium azide and add either 1µL of 1mg/mL BODIPY 493/503 in DMSO, 1 µL LipidSpot 610 (1000x) (Biotium) in DMSO, or 10µL of 200mg/mL (Sigma Aldrich) in acetone.

To determine the distribution of LDs and GGs in centrifuged embryos, *in-vivo* centrifugation was performed as described^20^, followed by fixation and simultaneous LD/GG detection as above. For analyzing follicles, ovaries were dissected from females maintained on yeast at 25°C overnight. Samples were then fixed with 4% formaldehyde in PBS for 15 min, washed in 1xPBS/0.1%Triton X-100, and simultaneously stained for LDs and GGs as above.

### Live imaging

For live imaging involving dye injections, a previously published procedure was followed^14^. In short, embryos were collected on apple juice plates for the desired time, hand-dechorionated, transferred to a coverslip with heptane glue, desiccated, and placed in Halocarbon oil 700. Embryos were then injected with BODIPY 493/503 (1mg/mL), LysoTracker Red (1mM) or LipidSpot 610 (1000x) and imaged on a Leica Sp5 confocal microscope.

For live imaging of Glycogenin-YFP and YV autofluorescence, embryos were collected on apple juice plates for the desired time, hand-dechorionated, transferred to a coverslip with heptane glue, covered with Halocarbon oil 27, and imaged. To improve signal, flies were kept in the dark, and light exposure during embryo preparation was kept to a minimum.

### TEM

Embryos were collected from 7-to 14-day-old flies, dechorionated in 3% sodium hypochlorite, and washed extensively with distilled water. Embryos were fixed in 4%paraformaldehyde/2%gluteradlehyde/PBS with an equal volume of heptane added. The vials were shaken then left on an agitator for 20 minutes. After fixation, embryos were washed extensively with 1×PBS/0.1% Triton X-100, then transferred onto a piece of double-sided tape, adhered, then submerged with 1×PBS/0.1% Triton X-100. The embryos were then gently hand rolled using fine forceps until the vitelline membrane was removed. Embryos were transferred to a small glass vial. The embryos were then fixed a second time with 4%paraformaldehyde/2%gluteradlehyde/PBS, excluding the heptane, for 30 minutes. Embryos were then washed three times with 0.2 M sucrose in 0.1M cacodylate buffer. They were washed an additional 3 times in 0.1 M sodium cacodylate before post fixation in 1% osmium tetroxide for 2 hours followed by uranyl acetate enhancement in 0.5% uranyl acetate overnight at 4 °C. Specimen were washed and then dehydrated in a graded ethanol series, transitioned to propylene oxide and embedded in an Epon/Araldite resin. Thin sections were stained with 0.3% lead citrate and imaged on a Hitachi 7650 transmission electron microscope using an 11 MP Gatan Erlanshen CCD camera. TEM work was conducted at the Electron and Cryo Microscopy Resource in the Center for Advanced Research Technologies at the University of Rochester.

### Quantification of TEMs

To quantify the association of LDs and GGs, LDs were manually identified based on their appearance and size (diameter of 0.3-0.75µm), while GGs were manually identified based on the staining pattern and diameter (2-7µm). The two structures were labeled associated if the distance between them was less than 30nm (∼2pixels).

### Particle Image Velocimetry

We performed PIV as described in^14^. The motion of acidic organelles and LDs was captured simultaneously in the same embryos. Embryos were collected, staged, mounted on a coverslip, and co-injected with BODIPY 493/503 and Lysotracker Red as described above. Per genotype, three embryos were imaged at 25°C. Timeseries were captured by confocal microscopy at a rate of 1 frame per 30 seconds, within a superficial plane of the embryo. The raw timeseries were then analyzed, finding the 8^th^ nuclear division by first finding the division where nuclei arrive at the periphery (the 9^th^ division), then going back one contraction cycle. 10 sequential frames were taken from this division starting at the period of the highest lipid droplet motion determined empirically. The signal from the embryos was then isolated from these 10 frames using a mask. These processed frames were then fed into a PIV algorithm based on OpenPIV, a python-based PIV implementation, generating 9 frame transitions per timeseries. The PIV analysis script used is available as supplemental data from our previous publication ^14^. The output vectors for each transition were then processed to remove any vectors which failed to meet or exceeded the empirically determined vector boundaries. Then directionality was removed, the vectors were averaged across the transition, pixels were converted to microns, and these averages were plotted on violin plots to show that pulses were being captured.

To perform PIV analysis for YVs, the same procedure was followed, with the exception that embryos were not injected, and embryos were illuminated with a 405nm laser and yolk autofluorescence was captured by collecting emission in a 410-500nm window.

### Statistics

All statistics were done using Graphpad Prism. P values were calculated using 2-tailed, unpaired Student’s T-tests. At least 3 embryos were used per genotype. For the PIV statistical comparisons 3 embryos were used per genotype contributing 9 transitions per embryo, thus n = 27, at least, per genotype.

The questions we sought to answer with statistical tests were “Are the Jabba LD:GG complexes flowing slower then wild-type singular LDs?”. Having received a positive result from this question, we next asked “Is the flow generally slower in Jabba than wild-type or is the diminished LD:GG speed due to complex formation?” for which we used acidic organelles and YVs as unclustered controls. These are binary questions seeking to determine if two populations of related numbers (flow speeds of organelles in each genotype) were different, thus we chose to use Student’s t-tests.

## Supporting information

Supplemental Figur 3

## Acknowledgements

We thank the Blooming Drosophila Stock Center, the Kyoto Stock Center, and Ramona Lattao for fly stocks. We thank Zehra Ali-Murthy and Thomas Kornberg (University of California San Francisco) who provided us with the protocol for fPAS staining. TEM analysis was performed by Chad Galloway and Karen Bentley in the Electron Microscopy Shared Resource Laboratory at the University of Rochester Medical Center. We are grateful to Fredrick Gootkind for initial analysis of the LD clustering phenotype in *Jabba* mutants and Ethan Fisher for his work on fPAS staining. This work was supported by National Institutes of Health grants F31 HD100127 (to M. D. K.) and R01 GM102155 (to M. A. W).

## Authorship contributions

M.D.K. conducted most of the experiments and analyzed the data, under guidance by M.A.W. M.D.K. and M.A.W. designed the overall project. M.R.J., J.M.T, and A.M. contributed experiments, technical advice, and conceptual discussions. M.R.J. performed initial analysis of GGs by fPAS staining and of GG-LD interactions in *Jabba* mutants. J.M.T. performed *in vivo* centrifugation experiments. A.M. performed initial electron microscopy characterization of *Jabba* mutants and advised on details and troubleshooting for the electron microscopy analysis included in the manuscript. M.D.K. and M.A.W. wrote the manuscript. All authors critically read the manuscript and provided feedback. M.D.K and M.A.W. contributed funding by National Institutes of Health grants F31 HD100127 (to M. D. K.) and R01 GM102155 (to M. A. W).

## Competing interests

The authors declare no competing interests.

## Materials and Correspondence

Correspondence and material requests should be addressed to Michael A. Welte.

## Supplementary Figures

Supplementary Figure 1. GG detection in in-vivo centrifuged syncytial embryos. A, A’) Glycogenin-YFP expressing embryo. A = bright field image; expected location of organelles indicated on the left, according to ^19^ and this study. The least dense fraction (the refractive lipid droplet cap) is at the top; the densest fractions (i.e., the refractive yolk vesicles and clear glycogen) are at the bottom. Scale bar = 50 µm. A’ = merged fluorescent image of the same embryo showing Glycogenin-YFP (yellow) at the very bottom and YV autofluorescence (blue) in the layer above). B,C) Centrifuged wild-type (B) and *Jabba* mutant (C) embryos and stained for glycogen (fPAS, red) and LDs (BODIPY, cyan). In the wild type, glycogen and LDs are present at opposite poles. In the *Jabba* embryo, LD and glycogen are co-mingled.

Supplementary Figure 2. Glycogen distribution at various embryonic stages. A) Glycogenin-YFP at different stages showing the clustering of glycogen granules and exclusion of glycogen from the pole plasm/pole cells (arrowhead). Left: YFP channel; right: corresponding brightfield image. B, C) fPAS staining of wild-type embryos. B) At the onset of germ band extension, fPAS signal is dimming and predominantly present the yolk cell. C) At the end of germ band extension (>7 hrs post fertilization), fPAS staining is undetectable. D) Glycogenin-YFP embryo during dorsal closure (∼12 hours post fertilization). YFP signal is very faint and present in the yolk cell. Signal appears brighter at the periphery due to optical sectioning. Images taken by confocal microscopy. E) Time course of fPAS staining in wild-type and *Jabba* embryos imaged using epifluorescence. The middle panels are at later germ band extension like that shown in panel C. We notice no differences between the genotypes. All scale bars = 100 µm.

Supplementary Figure 3. A,B) TEM images of newly laid wild-type (A) and *Jabba* mutant (B) embryos. GGs pseudo colored red; lipid droplets pseudo colored green. Scale bars = 2 µm. C) Embryos of different genotypes stained for LDs (BODIPY493/503, cyan) and glycogen (fPAS). In *Stur, klar, and LSD2/dPLIN2* null embryos, LDs are not associated with GGs. An embryo with reduced *Jabba* dosage displays partial association. D) Live imaging of stage 2 wild-type and *Jabba* embryos injected with BODIPY493/503 (cyan) to track LD motion. Arrowheads track selected LDs through time. The direction of the cytoplasmic flow is indicated by the black arrow. E) Stage 2 wild-type and *Jabba* embryos injected with LipidSpot 610 to label LDs (magenta)and co-imaged for YV autofluorescence (blue). In the *Jabba* embryo, the LDs rings are not around YVs. C-E) Scale bars = 10 µm; images recorded by confocal microscopy. F) cartoon showing the gross directions of the flowing cytoplasm during stages 1-3.

Supplementary Figure 4. Jabba prevents inappropriate interaction between LDs and GGs. A) TEM cross section of < 1hr old wild-type embryo. B) TEM cross section of a < 1-hour old *Jabba* embryo, note the LDs bound to GG and the lack of LDs free in the cytoplasm. Scale bars = 20 µm.

## Videos

Supplementary Video 1. Wild-type embryo injected with BODIPY493/503 (displayed in inverted greyscale) to label LDs. The embryo is imaged in a subcortical plane. The video starts in stage 1-2 (<1 hour post fertilization) and captures 30 minutes real time. Scale bar is 100 μm.

Supplementary Video 2. Wild-type embryo injected with BODIPY 493/503 to label LDs (shown in greyscale). The embryo is imaged at 40 μm below the subcortical plane. The video captures about 90minutes real time. Scale bar is 100 μm.

Supplementary Video 3. Glycogenin-YFP embryo with YFP in greyscale. The embryo is imaged in a subcortical plane. The video starts in stage 1-2 and captures 20 minutes real time. Scale bar is 100 μm.

Supplementary Video 4. Glycogenin-YFP embryo with YFP in greyscale. The embryo is imaged in a subcortical plane. The video starts in stage 1-2 (<1 hour post fertilization) and captures 2 hours real time. Scale bar is 100 μm.

Supplementary Video 5. Jabba embryo injected with BODIPY 493/503 (displayed in inverted greyscale) to label LDs. The embryo is imaged in a subcortical plane. The video starts in stage 1-2 (<1 hour post fertilization) and captures 30 minutes real time. Scale bar is 100 μm.

Supplementary Video 6. Jabba embryo injected with BODIPY 493/503 to label LDs (shown in greyscale). The embryo is imaged at 40 μm below the subcortical plane. The video captures about2 hours real time. Scale bar is 100 μm.

## Notes

### Competing Interest Statement

The authors have declared no competing interest.

### Summary of Updates

Supplemental files uploaded

