## Supplementary figures and images for "Drosophila embryos spatially sort their nutrient stores to facilitate their utilization"

### Supplemental Figur 3

Supplementary Figure 3

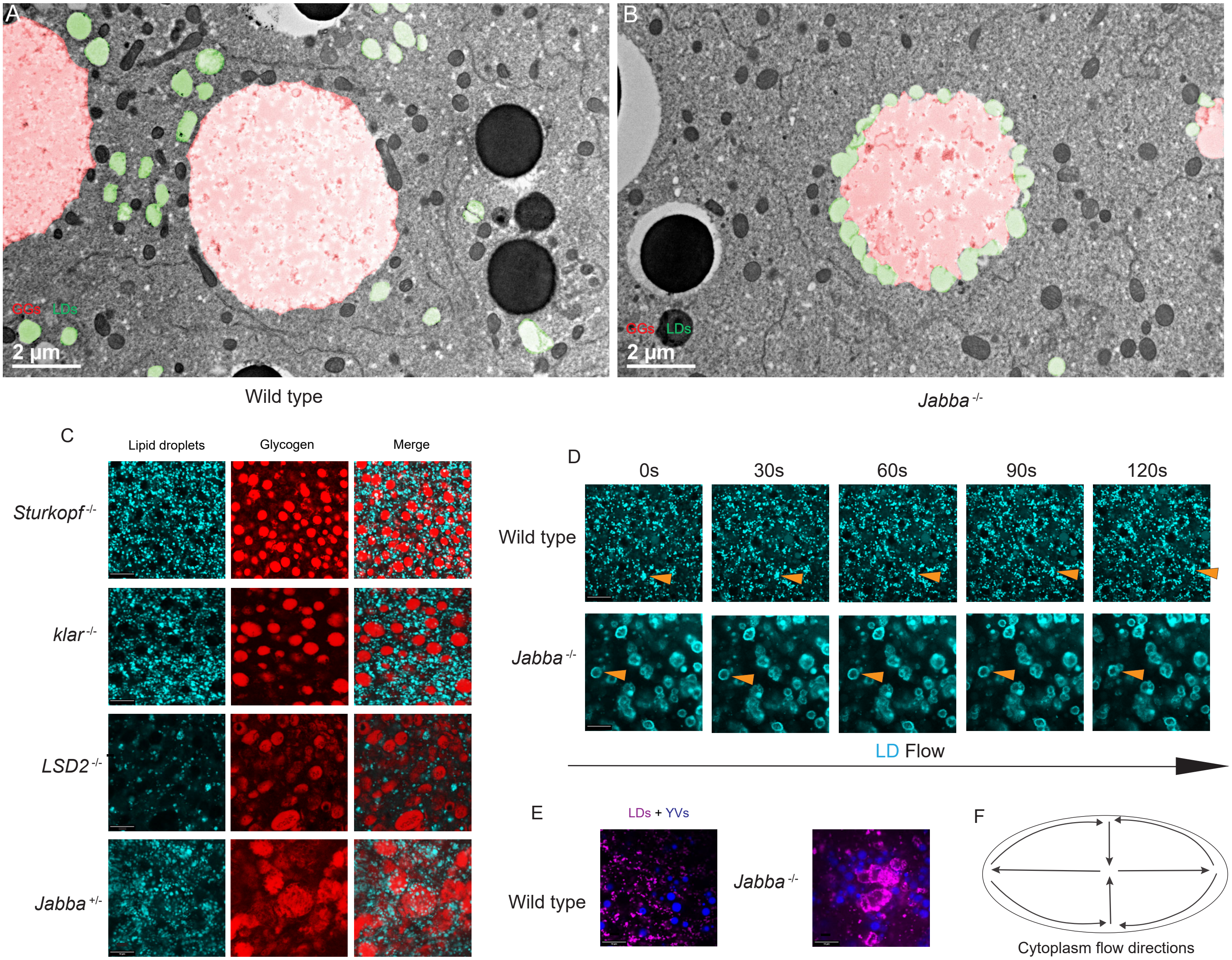

### Supplemental Figure 1

Supplementary Figure 1

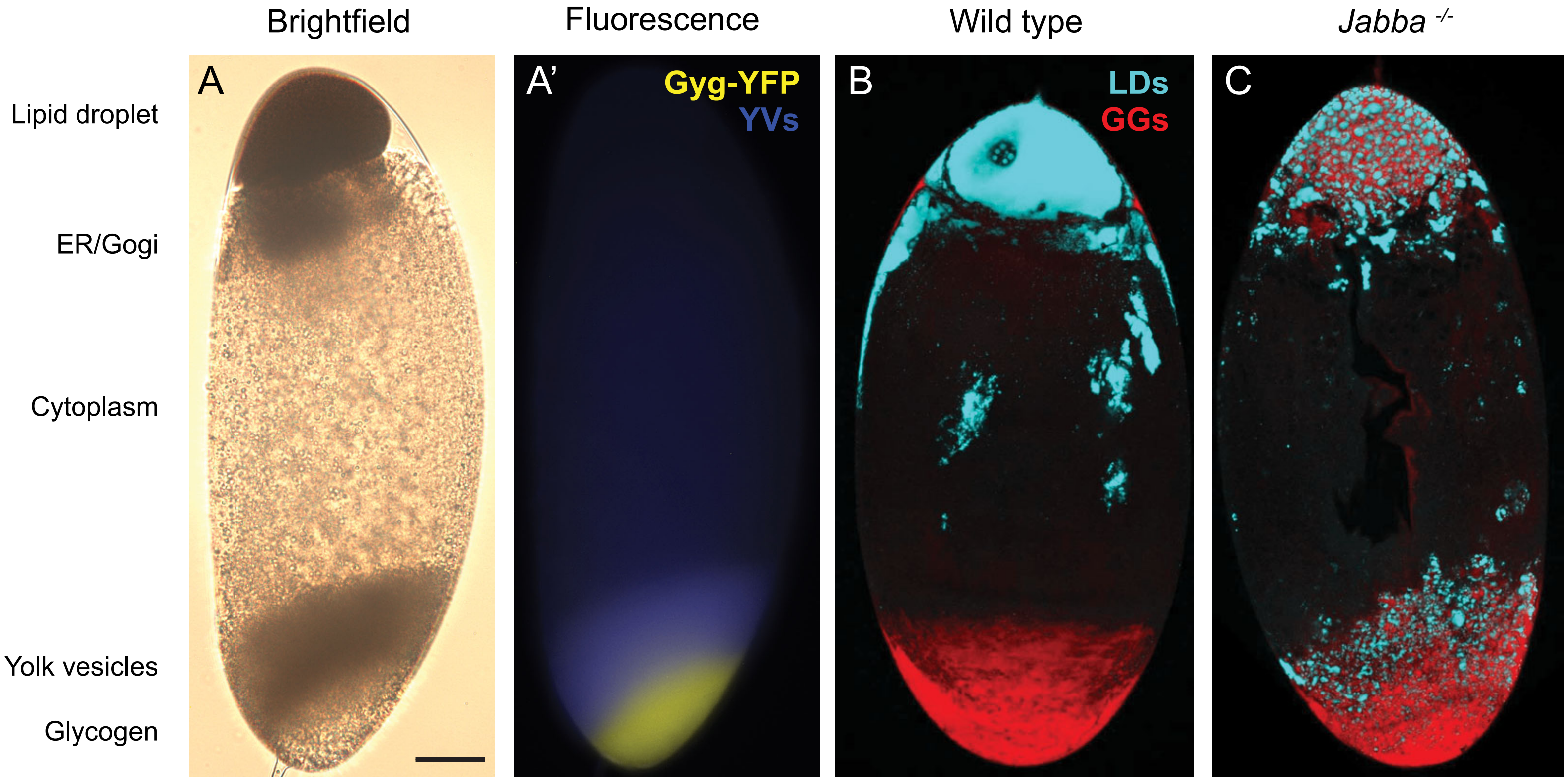

### Supplemental Figure 2

#### Supplementary Figure 2

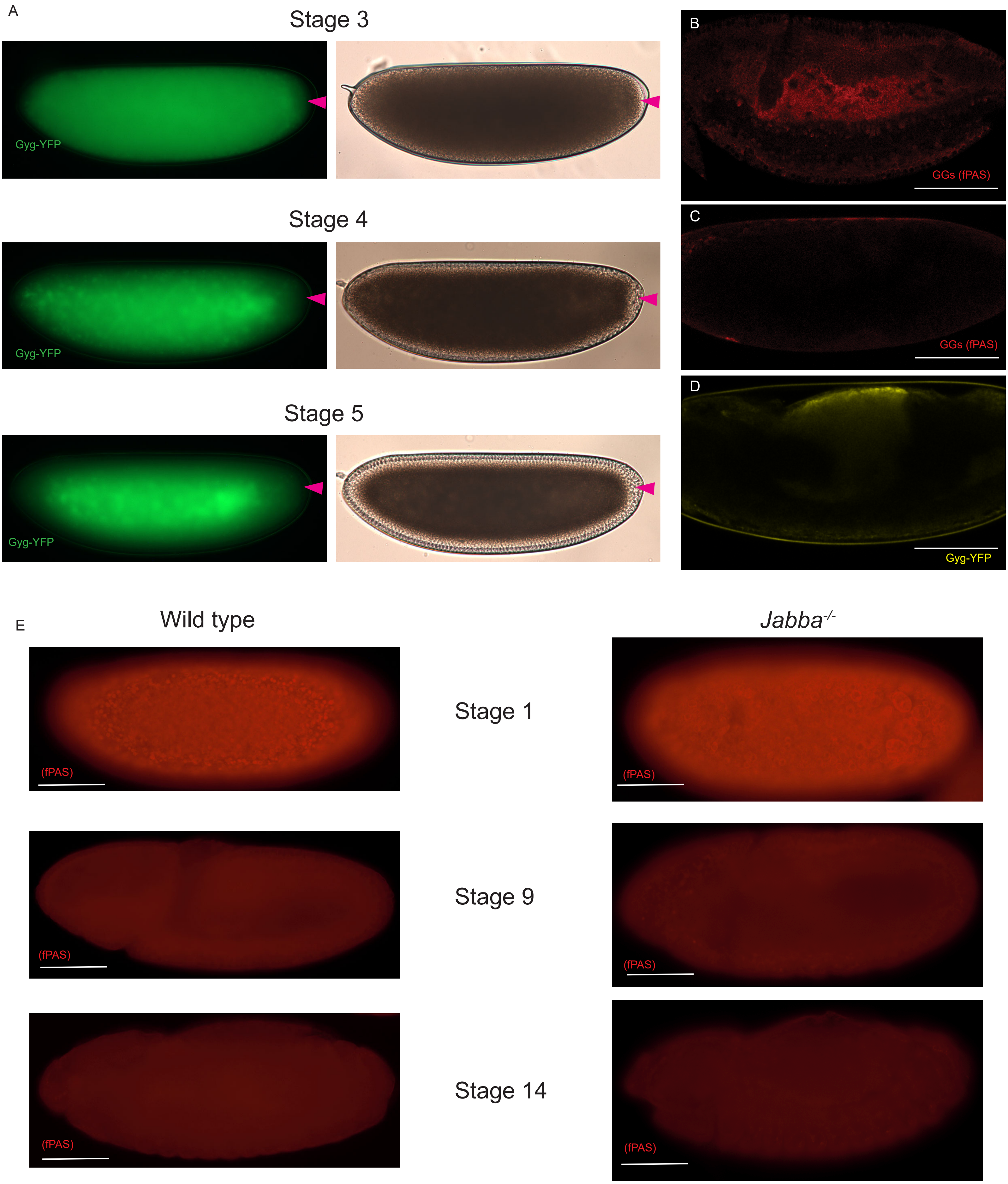

### Supplemental Figure 4

Supplementary Figure 4

Wild type

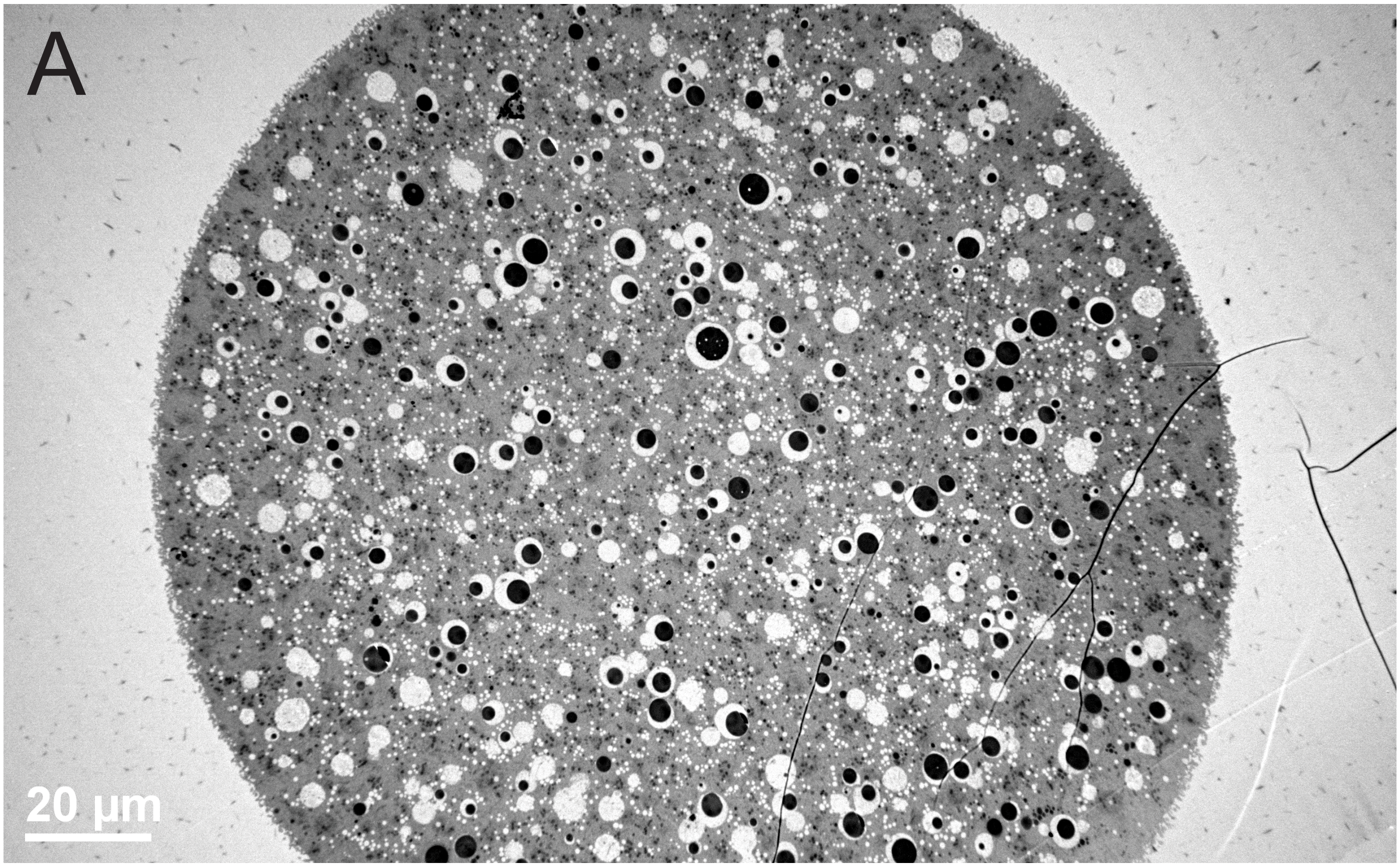


*Jabba* $^{-/-}$

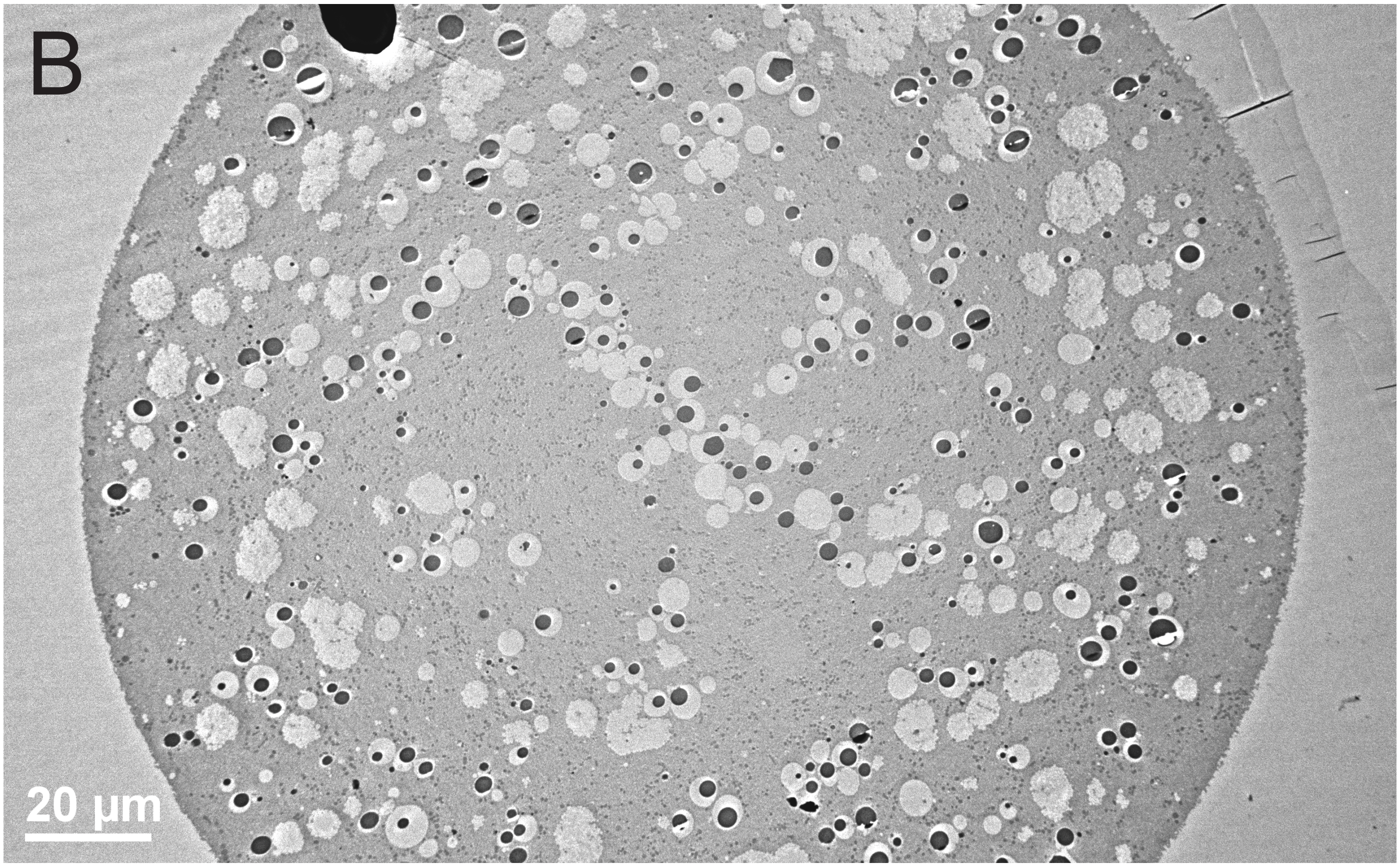
